# Integrative analysis of spatiotemporal transcriptomics delineates dynamic cell states in squamous tumorigenesis

**DOI:** 10.1101/2025.09.30.679409

**Authors:** Varsha Sreekanth, Timur Rusanov, Farah Yousef, Mark A. Sanborn, Su Yeon Yeon, Vijayalakshmi Ananthanarayanan, Olga Karginova, Jalees Rehman, Ameen A. Salahudeen

**Affiliations:** Department of Biochemistry and Molecular Genetics, University of Illinois, College of Medicine, Chicago, IL; Department of Biomedical Engineering, University of Illinois at Chicago, Chicago, IL; Department of Medicine, University of Illinois, College of Medicine, Chicago, IL; Department of Pathology, University of Illinois, College of Medicine, Chicago, IL; University of Illinois Cancer Center, Chicago, IL

## Abstract

Squamous cell cancers are responsible for 1 in 5 cancer deaths and survival improvements lag behind those seen in adenocarcinomas. This disparity is in large part due to the limited impact of immunotherapy due to therapeutic resistance, where less than ten percent of patients respond in early stage squamous cell carcinomas of the Head and Neck. Mechanisms that govern intrinsic resistance remain poorly understood and likely arise during the premalignant or dysplastic state. Here, we generated a dataset of murine and human oral squamous epithelia spanning the earliest premalignant stages through invasive carcinoma. Integrative analysis of single-cell and spatial transcriptomics data across the dysplasia to carcinoma continuum reveals early and sustained shifts in epithelial transcriptomes. Spatially informed cell-cell interaction analysis reveals dysplasia-specific upregulation of wound healing and immune remodeling programs in severe dysplasia and invasive carcinoma in a human sample of oral dysplasia. The presence of these transcriptomic programs within early dysplasia may account for the aggressive clinical presentations of squamous cancers including intrinsic therapeutic resistance.

## INTRODUCTION

Squamous cell cancers are responsible for 1 in 5 cancer deaths and survival improvements lag behind those seen in adenocarcinomas. In addition to landmark TCGA studies, recent literature has highlighted key differences in the biology of squamous carcinomas as compared to adenocarcinomas from the same tissues^1,2^, including immune and tumor oncogenic signatures associated with worsened outcomes^3,4^. Among these squamous cancers, squamous cell carcinomas of the head and neck are a leading cause of cancer-related death including HPV negative squamous cell carcinomas of the oral cavity (OSCC)^5^. Immunotherapy^6^ and other advances in the past decades have made minimal impact, and the survival rate in Early-Stage OSCC remains approximately 50%, as compared other cancers^5–7^.

Squamous cancers originate from epithelia possessing progenitor cells residing in the basement membrane adjacent or basal layer. These progenitors typically express TP63 and keratins including KRT5 and KRT14^8–11^ which accumulate DNA damage during exposure to carcinogens and chronic injury^12–14^. Studies of the premalignant epithelium suggest remodeling of stromal cells including fibroblasts and macrophages forms a microenvironment that promotes epithelial progression from early dysplasia to carcinoma, also known as tumor progression^15–19^. Early stage head and neck cancers and OSCC are characterized by high rates of recurrence, primarily due to intrinsic aggressive tumor biology that harbors intrinsic resistance to conventional chemotherapy and immunotherapies. This year marked the first advance in OSCC and head and neck cancer treatment, where the KEYNOTE-689 study^20,21^ demonstrated that in early-stage head and neck cancers, the anti-PD1 antibody pembrolizumab given before and after surgery (perioperative immunotherapy) significantly improved survival in early stage disease. However, the rates of immunotherapy response were less than 10%^21^ demonstrating that intrinsic resistance exists even in early-stage disease, and echoing the resistance observed in other neoadjuvant chemotherapy studies^7^. These results underscore the critical need to understand the mechanisms underlying this aggressive biology and improve outcomes by developing strategies targeting these mechanisms.

One approach to identifying the origins of this aggressive biology, as well as therapy resistance, is the use of unbiased single-cell sequencing of neoplastic lesions from the premalignant stage through invasive carcinoma. Such single-cell transcriptomic analysis would allow for an unbiased assessment of potential molecular pathways through the phases of squamous tumor progression. While there have been several studies that have analyzed the single-cell transcriptome in squamous cell carcinomas, there is a paucity of studies that perform a temporal analysis of dynamic transcriptomic shifts in various cell types from the earliest stages of premalignancy. Furthermore, premalignant lesions, especially in squamous epithelia, are defined by their disruption of normal histologic architecture and encroachment of the basement membrane. To inform our unbiased single cell transcriptomics analysis, we integrated these disease features via spatial transcriptomics.

Therefore, we generated a dataset of murine and human oral squamous epithelia spanning the earliest premalignant stages through invasive carcinoma. Integrative analysis of single-cell and spatial transcriptomics data across the continuum of dysplasia severity to carcinoma reveals early and sustained shifts in epithelial transcriptomes, with spatial analysis confirming localization to epithelial compartments. Additionally, we developed cellSpat, a computational framework that leverages spatial colocalization to uncover and rank the most biologically relevant ligand-receptor interactions. Spatially informed cell interaction analysis reveals dysplasia-specific upregulation of wound healing and immune remodeling programs, which were observed in severe dysplasia and invasive carcinoma using experimental models and in a human oral dysplasia sample.

## RESULTS

To study the dynamics of oral squamous epithelia during tumorigenesis, we utilized the murine 4-nitroquinoline-1-oxide (4-NQO) carcinogen induction model which recapitulates the genomic landscape of human OSCC^22^. Mice (n=42) were sequentially harvested in a schema enabling evaluation of n = 3 biological replicates for each sex over a 23-week interval for downstream histopathological grading and paired single-cell and transcriptomics sequencing (**Fig. 1A**). Tissues demonstrated a continuum of hyperplasia to dysplasia and invasive carcinoma toward the end of the experiment between week 20 and 23 (**Fig. 1B**). Mice that reached morbidity endpoints (e.g. weight loss) prior to the pre-set harvest time point were euthanized, mostly after week 20. Notably, we observed sex differences in dysplasia onset and progression (**Fig. S1**), with male mice experiencing higher grades at earlier timepoints and greater proportions of invasive grades overall compared to their female counterparts. Following harvests, tongues were graded and lesions scored by blinded histopathologic evaluation, with representative images in **Fig. 1C**. As expected, increasing architectural and cytological disturbances are observed with increasing dysplasia severity, mirroring the landscape of OSCC progression in humans. Pathology grading informed our approach for a paired scRNA-seq and spatial transcriptomics workflow. We devised a unique multiplexed approach leveraging tissue microarrays (TMAs), incorporating 71 tissues from the 118 collected throughout the time course of the study in 12 TMAs, increasing the scalability and throughput of our capture area based spatial transcriptomics assay (**Fig. 1D**). Additionally, we included 2 TMAs from untreated C57BL/6 mice to serve as experimental controls. TMAs maintained tissue integrity as observed from histopathological review of H&E slides. Following TMA construction, TMAs were used sequentially for spatial and scRNA-seq workflows.

**Figure 1.**
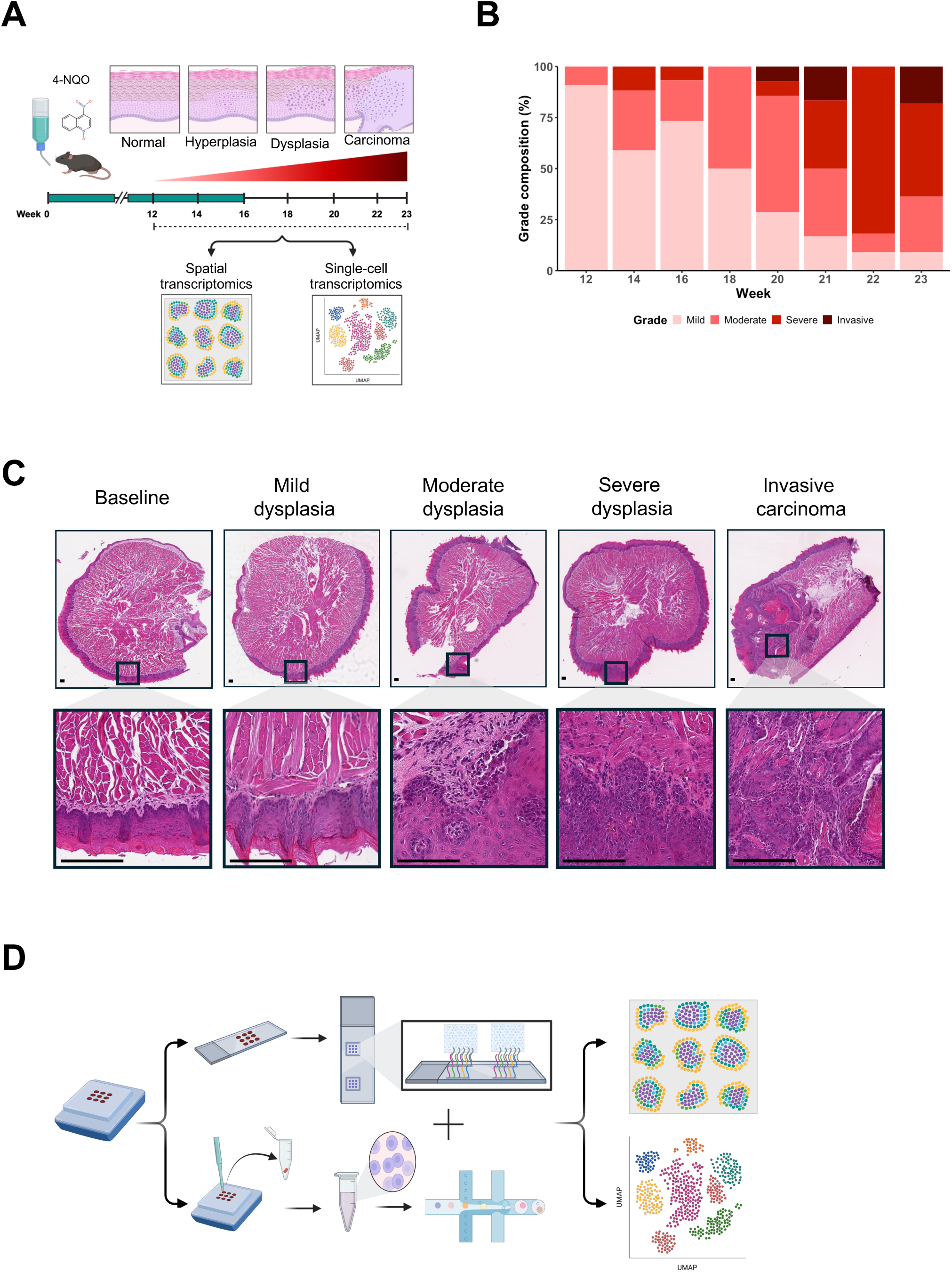
Modeling squamous tumorigenesis and integrative single cell and spatial analysis. (**A**) Schematic of murine 4-NQO induced carcinogenesis model. (**B**) Temporal progression of pathologist annotated dysplasia over 2 week intervals. (**C**) Representative H&E images of dysplasia grades, scale bar = 100 µm. (**D**) Schematic of paired scRNA-seq and spatial transcriptomics FFPE workflow and multiplexing. Mouse tongues collected throughout the study were embedded into tissue microarrays and subsequently processed for the capture area based 10X Genomics Visium spatial transcriptomics assay (top) then cells were isolated from the remaining individual tissues in the block for the 10X Genomics FLEX scRNA-seq assay (bottom). Data from both assays were subsequently integrated to enhance the resolution of spatial transcriptomics data and enable downstream bioinformatics analysis.

### Single cell transcriptomic analysis of oral carcinogenesis over time

Single-cell sequencing was performed on tissue samples across all dysplasia stages (**Fig. 1D**). High-quality cells were selected by preprocessing and doublet removal strategies as described in the Methods resulting in 150,344 sequenced cells meeting quality and doublet removal thresholds. scVI^23^ was used to generate a low-dimensional latent embedding and unsupervised clustering on the neighborhood graph from the latent space resulted in 10 major clusters annotated as unique cell-types using canonical marker gene sets. The global embedding was visualized with uniform manifold approximation and projection (UMAP) plots representing all the annotated cell types across mouse tongues (**Fig. 2A**) and was additionally stratified by sample and treatment (**Fig. 2B-C**, respectively). As expected, features plots demonstrated marker gene expressions within canonical lineages (**Fig. 2D**, **Fig. S2-3**). To analyze the shift in cell populations, we quantified the proportions of the different cell types across all the samples (**Fig. 2E**) which were simplified into biweekly intervals: weeks 12-14, weeks 16-18 and weeks 20 and beyond (**Fig. 2F**). Notably, the study ended at week 23 due to humane endpoints and we therefore combined this time point with the week 20-22 interval. We observed a trend of increased proportion of epithelial cells, consistent with hyperproliferation between baseline and increasing time intervals (**Fig. 2F**). Additionally, at the week 20-23 interval, the proportion of macrophages increased, the overall proportions of B-cells and T-cells were sustained, and the proportions of muscle cells, endothelial cells and fibroblasts decreased.

**Figure 2.**
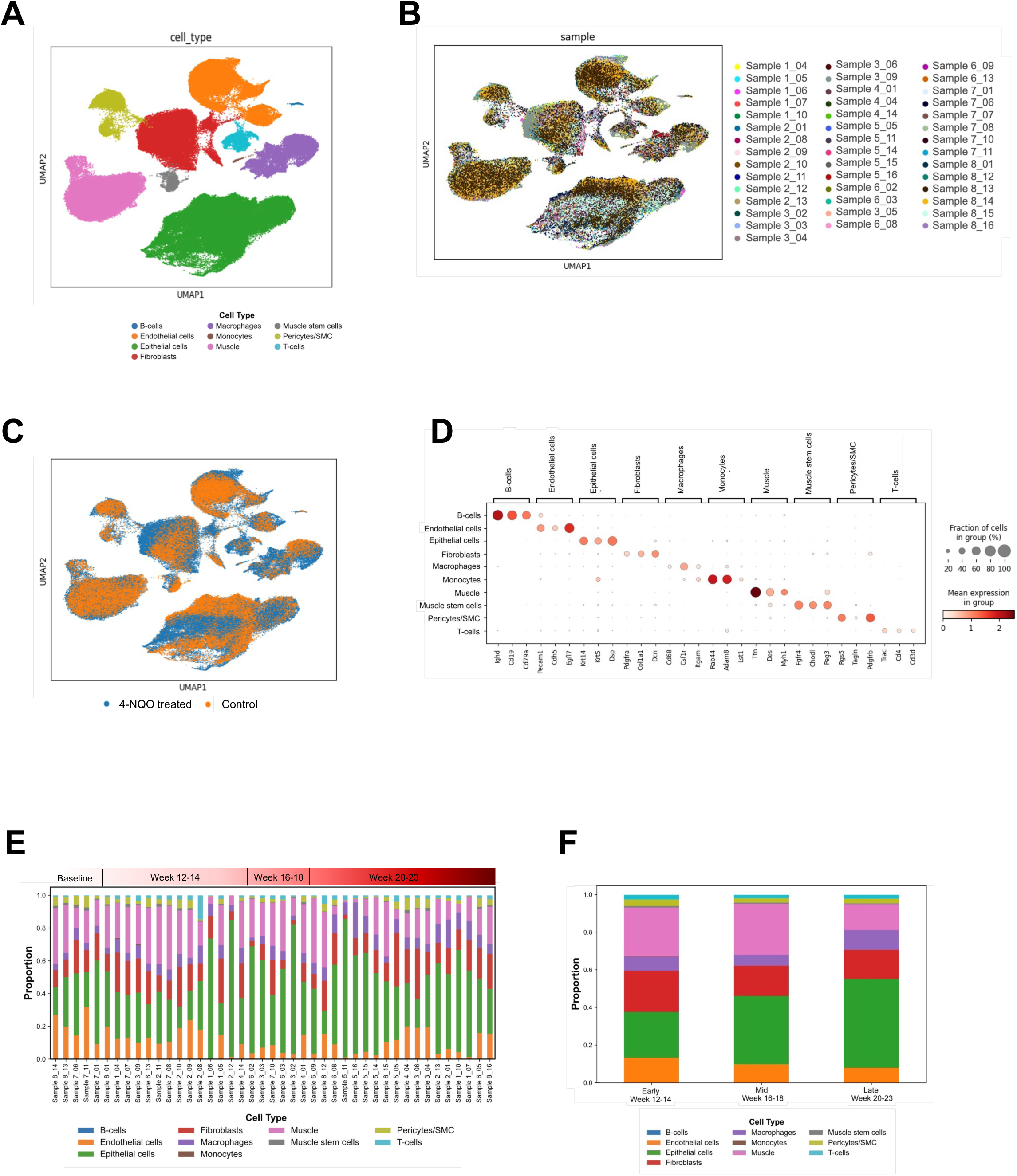
Murine oral lesion scRNA-seq cell clustering and annotation. (**A-C**) Low-dimensional UMAP visualization of single-cell sequenced tissue core samples clustered using the leiden algorithm and manually annotated cell types **(A)** using canonical marker gene expression and validation by feature plot analysis. Individual sample labels added as features over UMAP visualization, indicating sample distribution **(B**) and treatment conditions **(C**) of cells across clusters. **(D**) Dot plot of annotated cell-types, indicating mean expression per annotation group of marker genes associated with cell-type identity. **(E**) Cell-type proportions across samples. **(F**) Cell-type proportions across time intervals (weeks 12-14, 16-18, 20-23).

### Analysis of differential gene expression within the epithelium over time

Differentially expressed genes (DEGs) were determined via a pseudo-bulk expression matrix that was analyzed by DESeq2^24^ to characterize how the transcriptional profiles change throughout progressive disease stages in comparison to controls. In the epithelium, the total number of DEGs increased gradually with progression, with 1,256 DEGs for early lesions vs control to 1,442 DEGs for late lesions vs control, by the Wald test (**Supplemental Data S1-3**). Gene ontology (GO) enrichment showed upregulated DEGs corresponding to nuclear division, DNA replication, and cell differentiation pathways during early (week 12-14), mid (week 16-18), and late (week 20-23) phases of tumorigenesis (**Fig. 3A–C**). Downregulated DEGs showed GO enrichment for metabolic pathways and Wnt signaling (**Fig. 3D–F**). Key DEGs involved in nuclear division show continuous increases with disease progression (**Fig. 3G**). Conversely, genes involved in the negative regulation of Wnt signaling showed progressive downregulation over time (**Fig. 3H**).

**Figure 3.**
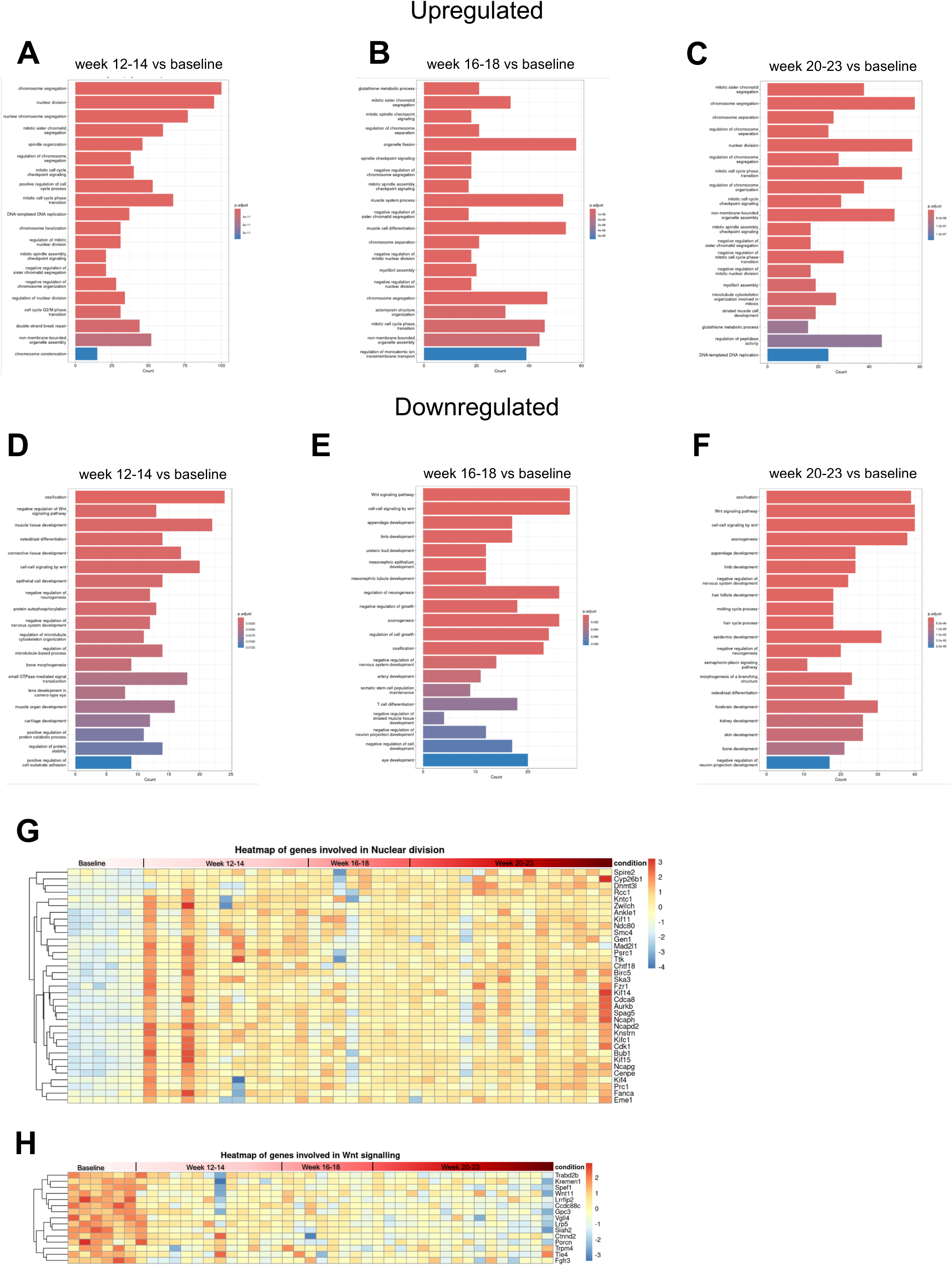
Differential expression analysis of epithelial cell scRNA-seq overtime. Upregulated (**A-C**) and downregulated (**D-F**) biological processes in the early (weeks 12-14) mid (weeks 16-18), and late (weeks 20-23) time point bins as predicted by gene ontology enrichment. (**G-H**) Heatmaps of differentially expressed gene expression involved in nuclear division (**G**) and Wnt signaling (**H**).

### Spatial transcriptomic analysis of oral cancer progression

We next performed spatial transcriptomics to analyze whether progression from dysplasia to oral cancer was associated with localized shifts in gene expression and cell profiles. To resolve the continuous variation of transcriptomes within each spatial spot, we deconvoluted cellular proportions using DestVI^25^ (**Fig. S4**), using the cell types identified in paired scRNA-seq data from the respective time points as the reference. Based on the distribution of cell-type proportions across spots, we applied cell-type–specific thresholds to assign a majority cell label, i.e., the dominant population within each spot.

In baseline samples (**Fig. 4A**), tissue architecture was preserved, with muscle cells occupying the central region, epithelial cells populating the mucosa, and minimal presence of infiltrated stromal and immune cells. However, in mild and moderate dysplasia samples, endothelial cells and fibroblasts began to infiltrate near the epithelial boundary (**Fig. 4B,C**). As lesions progressed to severe dysplasia and invasive carcinoma (**Fig. 4D,E**), the epithelial layer expanded, as indicated by the increased frequency of epithelial-majority spots. Fibroblast and macrophage accumulation proximal to the epithelium were observed in severe dysplasia, and were admixed with tumor cells in invasive carcinoma (**Fig. 4D,E**)

**Figure 4.**
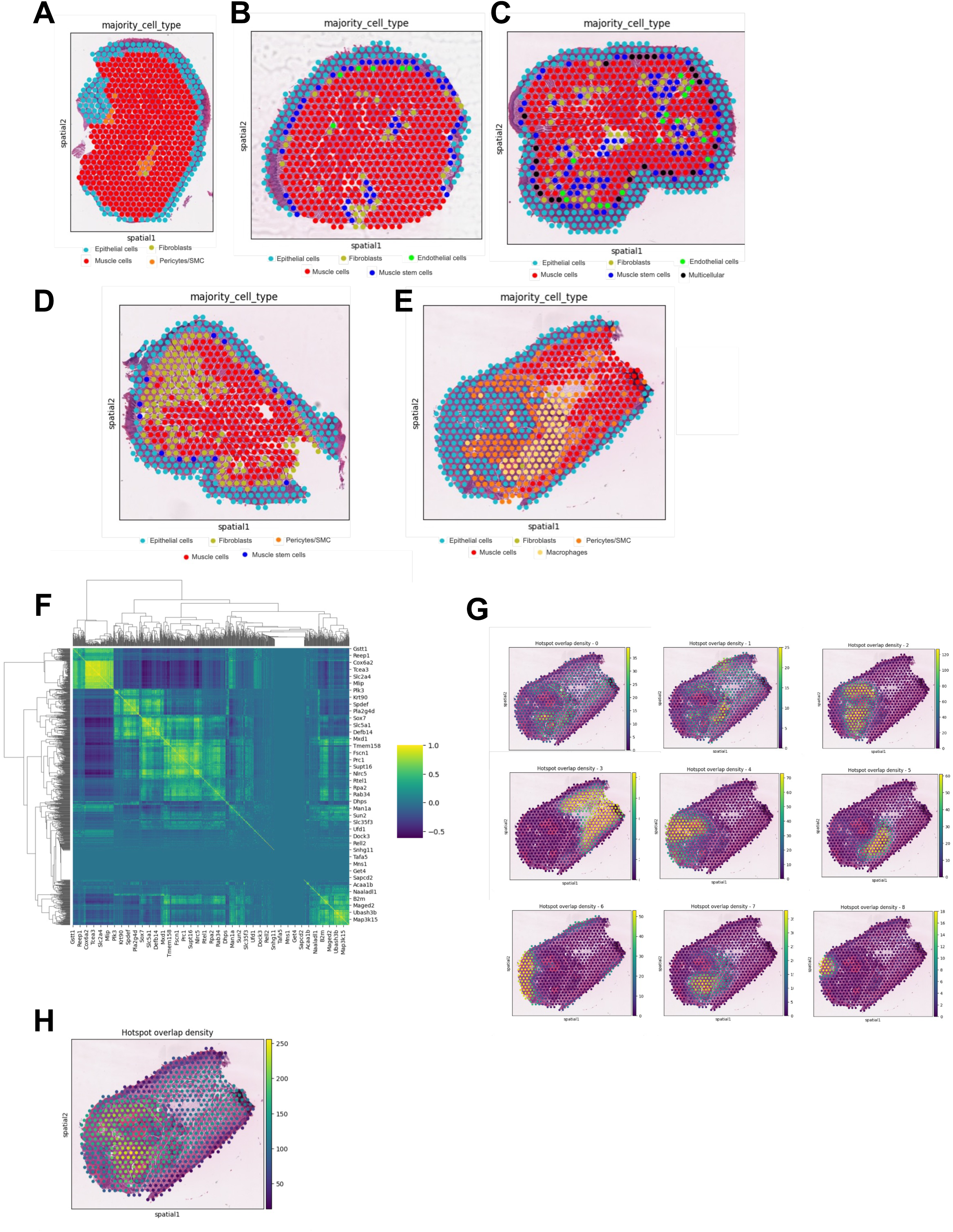
Spatial transcriptomic analysis of murine tongues. (**A-E**) Cell type proportions estimated following deconvolution with DestVI in normal, mild, moderate, severe, and invasive carcinoma respectively. (**F**) Heatmap of gene clusters based on spatial colocalization using Matthews correlation coefficient. (**G**) Hotspot density plots of genes present in individual gene clusters 0-8. (**H**) Overall hotspot densities of genes upregulated in invasive carcinoma.

### Spatial Differential Expression

To analyze spatial differential gene expression, we integrated paired scRNA-seq data to identify differentially expressed genes (DEGs) across temporal disease bins. Spatial hotspots of DEG expression were mapped using the Getis-Ord Gi* statistic^26^, and unsupervised clustering was performed based on the Matthews correlation coefficient^27^ to find clusters of spatially co-expressed genes. As a representative case, we focused on the invasive carcinoma sample, which was enriched for genes upregulated at the late time point and identified nine highly correlated Louvain clusters (**Fig. 4F**).

Hotspot density plots of DEGs within these clusters revealed co-localization patterns centered on the invasive lesion (**Fig. 4G**), with overall hotspot densities of late upregulated genes also localizing to this region (**Fig. 4H**). GO enrichment analyses of scRNA-seq data all converged on pathways implicated in nuclear division, DNA replication, and cell cycle progression, processes that are hallmarks of proliferative expansion in aggressive tumors. Collectively, this demonstrates that our hotspot approach provides an unbiased and zone-agnostic strategy to uncover spatially organized transcriptional programs, thereby correlating spatial gene expression heterogeneity to the underlying biology governing tumor growth.

### CellChat Analysis

Tumor progression in OSCC has been associated with remodeling of the microenvironment in pre-malignant lesions as they progress toward invasive carcinoma with shifts in the cellular communication landscape during tumor progression. CellChat analysis of scRNA-seq data revealed consistent epithelial–fibroblast interactions across all time intervals (**Fig. 5A–D**), highlighting the persistence of signaling between tumor epithelium and stromal fibroblasts, which has been known in literature to promote extracellular matrix remodeling and tumor invasion^28^. In contrast, monocyte and B-cell interactions progressively decreased over time. Among the interactions during week 12-14, epithelial fibroblast signaling dominated, with each cell type engaging in over 100 unique interactions (**Fig. 5E**). At week 16-18 time points, muscle cell–derived interactions began to wane, while epithelial, fibroblast, and endothelial cell interactions peaked, consistent with the role of fibroblast activation and stromal remodeling in dysplasia progression (**Fig. 5E**). At week 20 and beyond, macrophage– epithelial interactions became prominent suggesting the emergence of tumor associated and tumor promoting macrophages (**Fig. 5E**). Overall, the number of predicted ligand–receptor interactions diminished over time, decreasing from 403 in the early stage to only 50 by the late stage (**Fig. 5F**).

**Figure 5.**
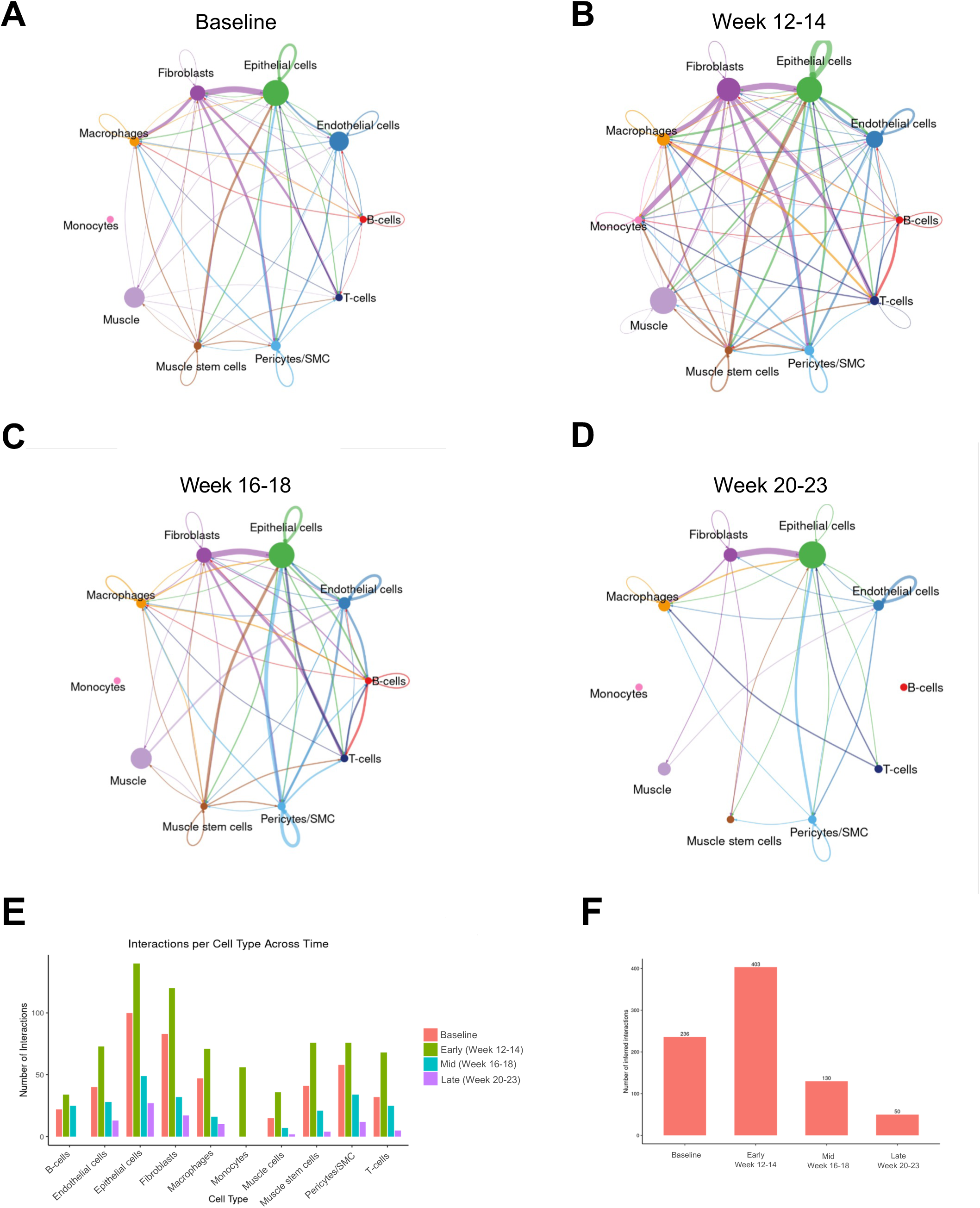
CellChat analysis of inferred cell-cell communication. (**A-D**) Circle plots indicating number of interactions inferred by CellChat between distinct cell-types from single-cell sequencing of tongue samples. Interactions from baseline (normal untreated tissues), early (weeks 12-14), mid (weeks 16-18) and late (weeks 20-23) samples plotted as **A, B, C,** and **D** respectively. (**E**) Overall interactions per cell-type across the tumorigenesis time-course stages. (**F**) Total number of inferred interactions across time-course stages.

### Spatially informed cell-cell communication prediction

CellChat predicts a large number of potential cellular interactions from scRNA-seq data alone, but predicted putative interactions of isolated cells that are sequenced in scRNA-seq analysis may not actually occur, unless receptors and ligands are spatially colocalized. To enhance the robustness and rigor of cell-cell communication prediction we developed cellSpat (https://pypi.org/project/cellspat/), a Python-based framework that integrates spatial transcriptomics with single-cell RNA-seq. CellSpat leverages the transcriptionally inferred interaction networks chrome CellChatv2 and ranks ligand–receptor (L-R) pairs based on their degree of spatial co-localization in a statistical framework of a hypergeometric distribution. Using cellSpat, we found that App–Cd74 interactions consistently emerged as statistically significant over the time course of tumor progression, as well as Fgf7-Fgfr1 interactions (**Fig. 6A–D**). App-Cd74 is part of an axis of the prototypic macrophage inhibitory factor (MIF) and its receptor, Cd74, and is integral to priming macrophages in wound healing, similar to the CCN1 integrin axis^29,30^. APP-CD74 has also been implicated in disease progression in various solid tumors as well as hematologic malignancies^31–33^. To further resolve where these signals concentrate, we applied cellSpat’s spatial autocorrelation module, which identified hotspots (clusters of ligand and receptor co-expressing spots that co-occur more frequently than expected by random chance) of L-R co-localization. App– Cd74 hotspots localized predominantly at the epithelial boundary (**Fig. 6E–H**). Feature plots of CD74, its coreceptors, and ligands, including macrophage inhibitory factor (MIF) and its related gene DDT, showed greater colocalization in areas of severe dysplasia and foci of invasive carcinoma (**Fig. 7A-B, D-E**) as compared to the CCN1-integrin axis associated with wound healing (**Fig. 7C,F**).

**Figure 6.**
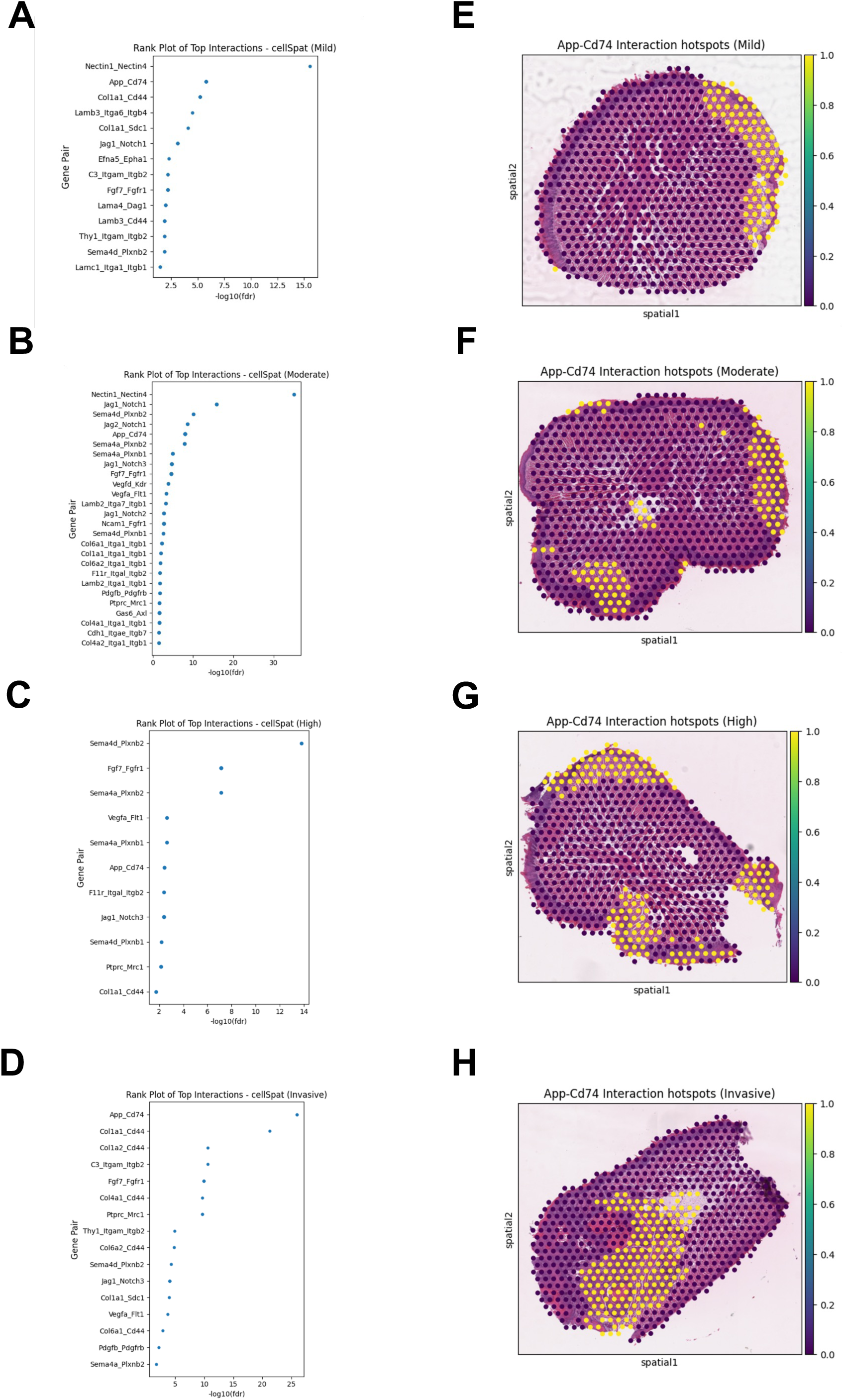
Spatially informed cell-cell communication analysis using CellSpat. (**A-D**) Top ranked cellSpat ligand-receptor interactions in representative mild, moderate, severe, and invasive carcinoma samples. (**E-H**) cellSpat predicted hotspots of the App-Cd74 interactions demonstrate enrichment within dysplastic epithelia.

**Figure 7.**
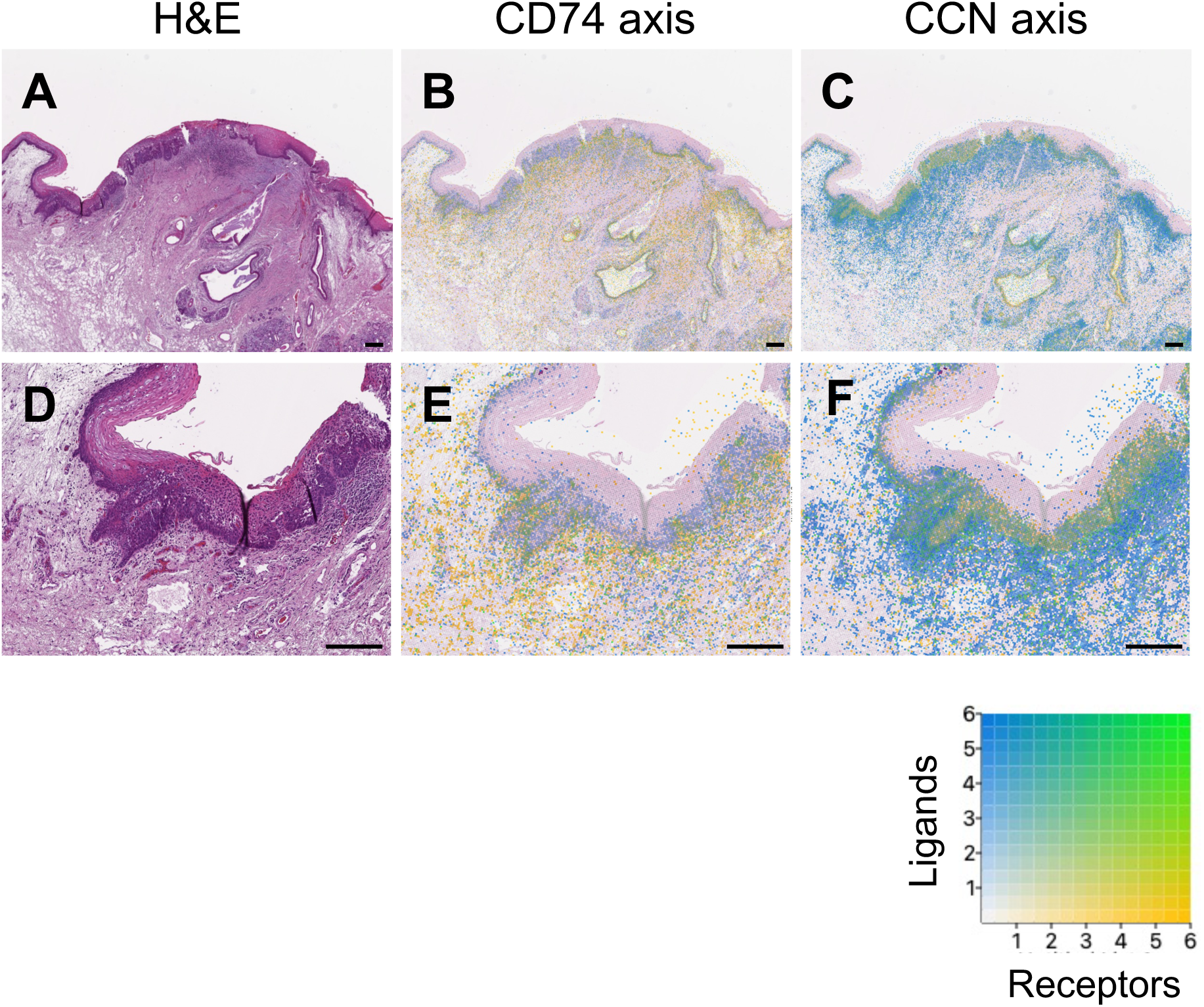
Spatial transcriptomics analysis of human oral dysplasia. (**A**) H&E of severe squamous dysplasia with focal invasive carcinoma, scale bar = 200 μm. (**B**) Colocalization of summed log-normalized counts of ligands (DDT, MIF, APP) and CD74 and coreceptors (CD74, CD44, CXCR2, CXCR4). (**C**) Colocalization of summed log-normalized counts of CCN1 family ligands (CCN1-6) and receptors (ITGA2, ITGA5, ITGA6, ITGAV, ITGAM, ITGB1-3, ITGB5). (**D-F**) High magnification of A-C, respectively.

## DISCUSSION

Squamous cell cancers including head and neck cancers have shown benefit with immune checkpoint blockade, yet clinically significant response rates in the neoadjuvant setting only occurs in a minority of patients. Here, we observed remodeling of the epithelia away from homeostatic renewal and differentiation pathways and toward wound healing and proliferation pathways. This remodeling is associated with ligand-receptor interactions, which were deciphered in silico. CellChat infers potential ligand–receptor interactions solely from single cell transcriptional profiles, and its successor CellChatv2^34^ allows for the integration of spatial transcriptomic data by analyzing of interactions between spatial locations. However, this approach is limited by the heterogeneity of cell types in any given spot. Therefore, we developed cellSpat, which predicts cell–cell interactions by ranking ligand–receptor pairs inferred from single cell transcriptomic data and then assesses spatial co-localization of those interactions. Through this framework that integrates spatial transcriptomics with single cell transcriptomics, we observed a marked increase in autocorrelated App–Cd74 signaling during the progression of oral cancer in the experimental murine model. In human, we also observed colocalization of CD74 ligands including APP, MIF, and DDT and CD74 within focal areas of an oral dysplasia and invasive carcinoma sample. MIF-CD74 has been associated with tumor progression and CD74 has been studied in the context of wound healing^30,35^ where ligands MIF and DDT can engage CD74 along with co-receptors or alternative receptors CXCR2 and CXCR4^31,36^. With respect to squamous cancers, MIF upregulation has been associated with worse overall survival in head and neck cancers^37–39^ and upregulation of its receptor, CD74 has been shown to correlate with tumor progression in oral squamous cancer^40,41^.

Another feature of tumor progression that was evident in our analysis was the shift of stromal and immune populations over time. Deconvolution of spatial data identifies endothelial and fibroblast residing at the epithelial boundary in mild/moderate dysplasia, followed by fibroblast and macrophage accumulation adjacent to or admixed with tumor epithelium in high-grade lesions and invasive carcinoma. CellChat analyses are consistent with this trajectory: epithelial–fibroblast signaling persists across time, while monocyte- and B cell– mediated interactions wane and macrophage–epithelial interactions become prominent at late stages including the putative App-Cd74 as above. The overall trend suggests that over the course of tumor progression, heterotypic cell interactions converge onto pro-tumorigenic circuits consisting of epithelial-fibroblast and epithelial-macrophage interactions. These findings are in line epithelial driven remodeling of the stroma^28,42,43^ and will be require further mechanistic experiments for validation.

## LIMITATIONS

This study has several limitations. The spatial transcriptomics cellular resolution was limited due to spot size, where deconvolution could only infer precise cell type composition, requiring single cell resolution confirmation with evaluation of corresponding protein expressions. While the murine model has been shown to recapitulate key genomic features of human OSCC, it may not fully capture etiologic diversity or the full immune landscape, requiring a more comprehensive data set of human dysplasia data to support our observations in the mouse.

In summary, integrated analysis of spatiotemporal data delineates early epithelial and microenvironmental programs that persist and increase during oral squamous tumor progression, thereby providing a framework to explain aggressive tumor biology and intrinsic resistance in squamous cancers. Targeting these circuits alongside immune checkpoint blockade may be a strategy to improve response rates in the neoadjuvant setting, especially in OSCC.

## ACKNOWLEDGEMENTS

We wish to thank core facility members and leadership at the University of Illinois Cancer Center, University of Illinois Chicago Research Resources Center, Biologic Resource Laboratory, and the University of Illinois Urbana-Champaign’s Roy J. Carver Biotechnology Center, DNA Services core. This study was supported by NIH grants K08DE027730(A.A.S.), F31AG090005(M.A.S), R33CA258012(J.R.), and R01HL163978(J.R.).

## METHODS

### Key Resources Table

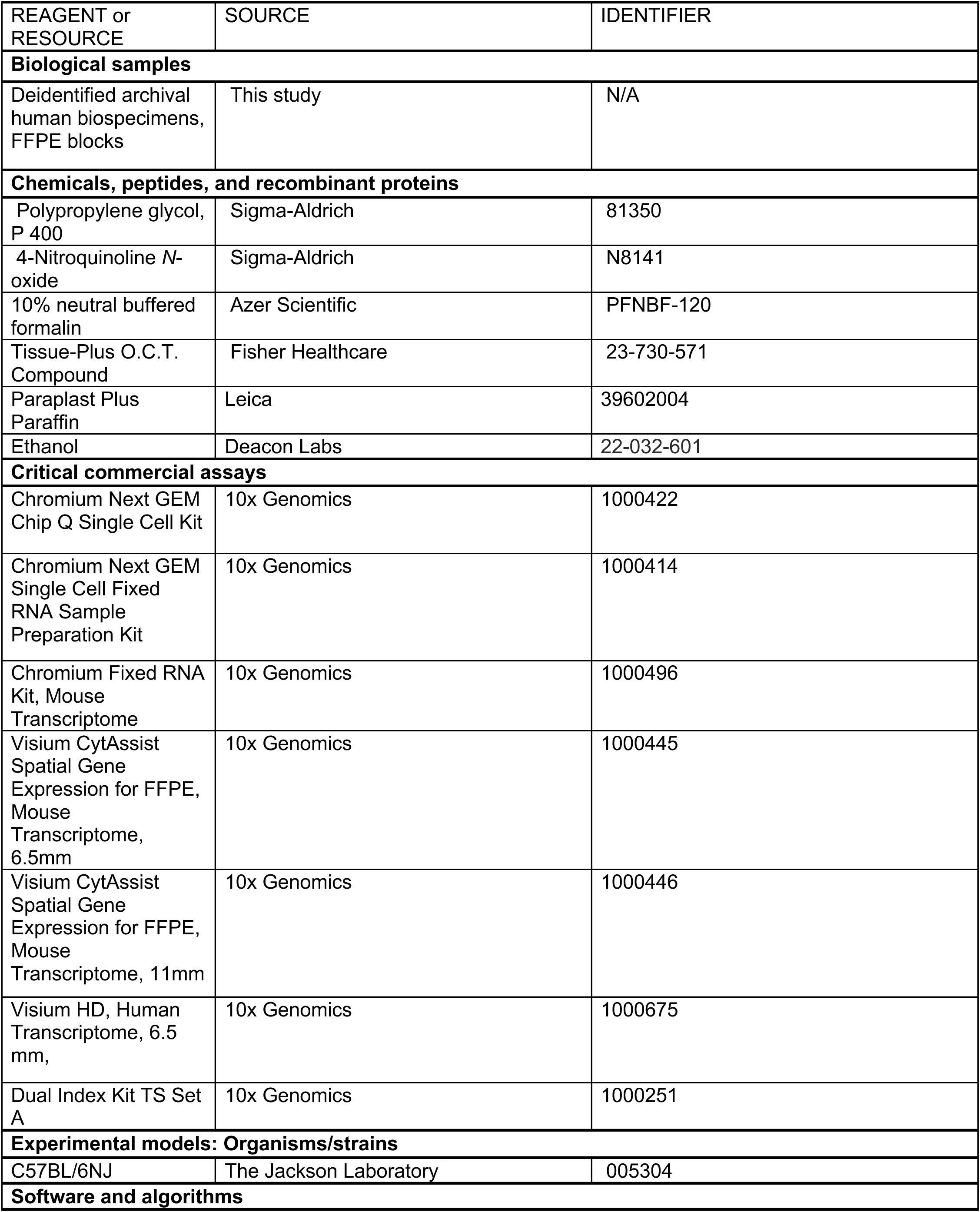
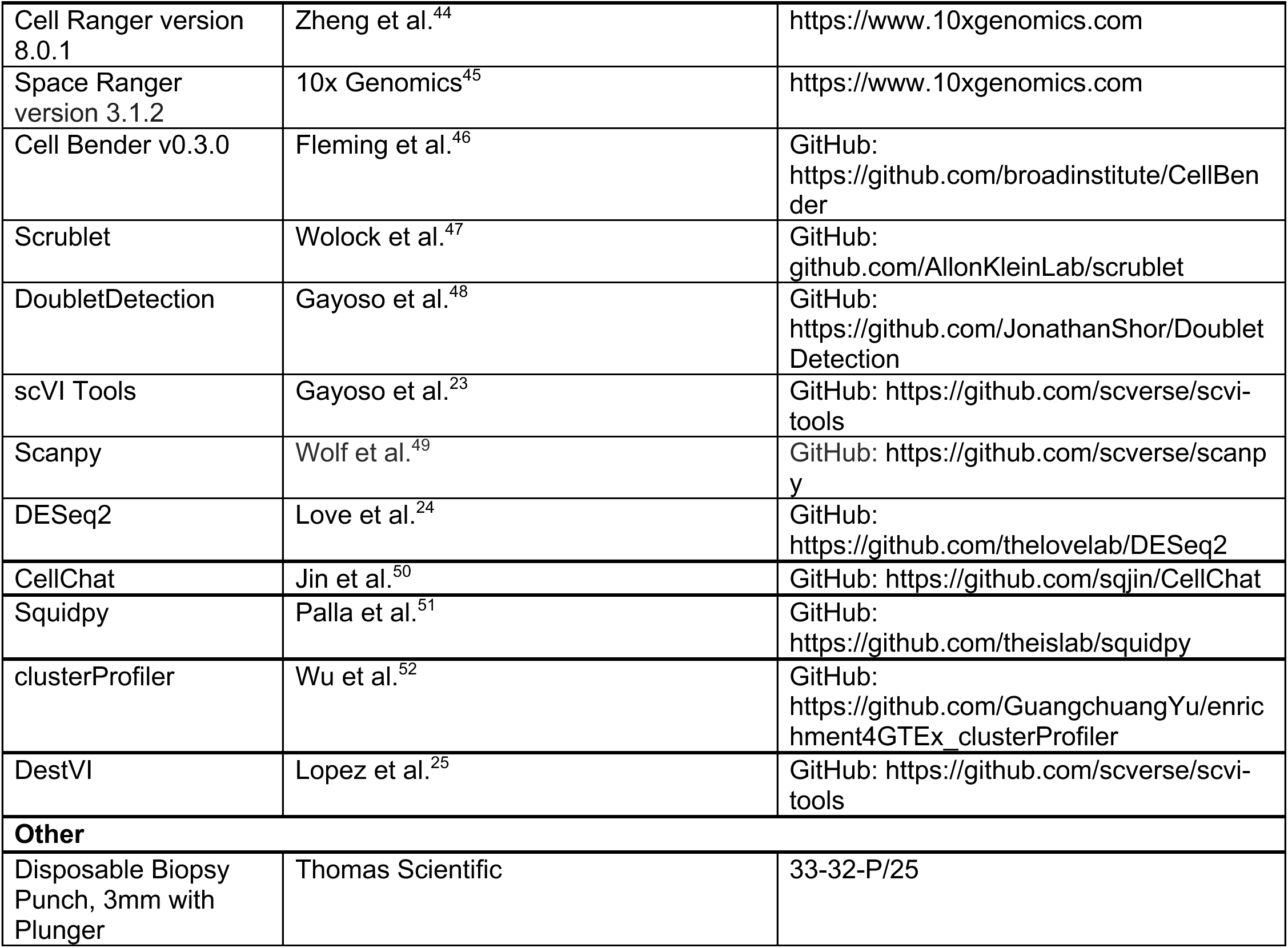

### Experimental models and samples

This study was carried out under protocols approved by the University of Illinois Chicago IACUC (AWA #D16-00290-A3460-01) and Institutional Review Board (FWA #00000083).

### Murine 4-nitroquinoline-1-oxide carcinogen induction model

Murine oral squamous cell carcinomas were induced in C57BL/6 mice using the 4-nitroquinoline-1-oxide drinking-water model as previously described^22^. Six- to eight-week-old animals of both sexes (n=42) were exposed to 4-NQO (100 μg/mL) in drinking water for 16 weeks and then returned to regular water until tissue harvest. 4-NQO stock was prepared in 100% polypropylene glycol and diluted into drinking water to a final 0.2% vehicle; solutions were refreshed weekly. Mice were housed under standard conditions, weighed and observed weekly, and were sacrificed at humane endpoints when indicated. Beginning at week 12 of induction, sex-balanced cohorts were sacrificed at two-week intervals to collect tongues for analysis. Following CO₂ euthanasia, whole tongues anterior to the circumvallate papillae were excised and cut coronally into 2-mm segments (4–6 per animal). Alternating segments were preserved either in a) snap frozen and embedded in OCT blocks or b) Formalin Fixed Paraffin Embedded blocks (FFPE) for downstream assays.

### Histological evaluation and scoring

Histopathological assessment of the tongue specimens was performed by a board-certified pathologist who was blinded to the experimental groups. The evaluation was conducted in accordance with the World Health Organization (WHO) 2017 criteria for the diagnosis of epithelial dysplasia^53^. These criteria encompass both architectural and cytological features. Architectural features assessed included basal cell hyperplasia, sharp demarcation from adjacent inflamed or normal mucosa, presence of abnormal superficial mitotic figures, and loss of epithelial maturation. Cytological features evaluated included variation in cellular size and shape, increased nuclear-to-cytoplasmic ratio, elevated mitotic activity, and presence of atypical mitotic figures. Based on these criteria, lesions were graded using a three-tier classification system as follows: mild dysplasia, moderate dysplasia, and severe dysplasia. Cases exhibiting invasion beyond the basement membrane were classified as invasive carcinoma.

### Tissue microarray construction

Tissue microarrays (TMAs) were constructed to scale the number of samples per spatial transcriptomics capture area. FFPE mouse tongue slices (n = 118) representing tissue collected from all treated animals were sectioned. H&E stained sections were reviewed by a pathologist, and 71 tissue slices were selected for TMA construction, capturing representative dysplasia grades by time point and by sex. Tongue sections from mice at baseline were included as experimental controls. To construct the TMAs, whole tongue circular cores (3 mm diameter) were extracted from the donor blocks with a biopsy punch (Thomas Scientific) and re-embedded according to a pre-configured map into a new paraffin block. TMAs were designed with either 2 x 2 or 3 x 3 tongue tissues to fit into Visium standard definition capture area (6.5 x 6.5 mm or 11 x 11 mm). A total of 12 TMAs (71 tissues) from 4-NQO induced carcinogenic mouse model were produced, in addition to 2 TMAs (8 tissues) constructed with tissues from untreated baseline controls. Following TMA construction, FFPE blocks were sectioned and H&E-stained to verify tissue integrity, orientation, and grades prior to spatial and scRNA-seq workflows.

### Spatial transcriptomics

#### Visium Assay

Mouse FFPE TMA samples were sectioned, H&E stained, imaged, processed and sequenced according to 10X Genomics Visium CytAssist Spatial Gene Expression User Guides for FFPE (CG000495). Mouse v2 probes were used for probe hybridization. Human archival FFPE samples were processed using 10X Genomics Visium HD Spatial Gene Expression User Guides (CG000685). Space Ranger version 3.1.2 was used for data preprocessing. Remaining tissue in the FFPE blocks after the spatial assay were used for cell isolations followed by scRNA-seq performed with 10X Genomics Chromium Fixed RNA Gene Expression Profiling (FLEX).

#### Visium HD Assay

Human Oral Tissue FFPE blocks were archived under protocols approved by the Institutional Review Board at University of Illinois at Chicago. All diagnoses were verified by a board-certified pathologist. FFPE blocks were sectioned and H&E stained to identify an area of dysplasia and evaluate its degree. 2×25 μm scrolls and 5 μm unstained consecutive sections were produced for 10X Genomics FLEX scRNA-seq and Visium HD spatial assays, respectively, followed by an additional 5um H&E stained section used as quality control. Samples were further processed according to 10X Genomics user guidelines to immediately proceed with downstream assays.

#### Cell Isolations and scRNA-sequencing

Mouse tissues remaining in the FFPE blocks after the spatial assay were individually cored for cell isolations. Human tissue scrolls used for cell isolations were from the same pathologist-reviewed archival FFPE blocks utilized for spatial transcriptomics assay. Workflows with human and mouse samples were conducted separately. Cells were isolated using 10X Genomics protocols for Chromium Fixed RNA profiling (CG000632). Tissues were digested for 45-60 minutes. Cells were stored according to the manufacturer’s recommendations and were defrosted and washed on the day of probe hybridization to proceed with the workflow. Individually barcoded samples from the same species were multiplexed and pooled for GEM generation, 5-10 samples per gel bead. Libraries were prepared according to 10X Genomics User Guides for multiplexed samples (CG000527). Cell Ranger version 8.0.1 was used for data preprocessing.

#### Data Preprocessing

Pre-processing scRNAseq data – FLEX-Libraries were sequenced with Illumina NovaSeqX chemistry targeting an average of 10,000-20,000 reads per cell according to 10X Genomics recommendations. 10x Genomics Cell Ranger pipeline (version 8.0.1) using the mm10 genome reference obtained from 10x Genomics demonstrated an average library read depth of 10,000 reads per cell for each barcoded library pool. Barcoded libraries that did not meet a minimum 300 median genes per cell were removed from downstream processing.

Dead cell and doublet removal – High quality cells were identified using a combination of ambient RNA removal, CellRanger cell annotations, and automated and manual doublet removal. Ambient RNA was removed using CellBender v0.3.0^46^ with a learning rate of 1e^-4^ with a default of 150 epochs and cuda enabled. Cells not identified by CellRanger were removed from CellBender-corrected matrices. Cells with fewer than 200 detected genes and greater than 15% mitochondrial gene expression were further excluded. A median absolute deviation (MAD) filter was then used to discard cells whose total UMI counts, number of detected genes, proportion of top 20 most highly expressed genes, and proportion of mitochondrial transcripts that deviated more than *k* MADs from the median (usually 5 MADs for library size and gene count, and 3 MADs for mitochondrial content). For doublet removal, Scrublet^47^ (from scanpy v1.10.3) was first utilized with a 5% expected doublet rate. In parallel, DoubletDetection^48^ (v4.2) was used to compute doublet scores and binary doublet calls. The union of cells flagged as doublets were then removed from the dataset. Markers of canonical cell lineage (**Figures S2-3**) were then manually selected for pruning of Leiden clusters (see below) of putative doublets. A binary matrix was generated to represent marker gene expression (raw counts > 1 in a given cell). Clusters were then pruned if the proportion of cells co-expressing lineage markers were > 0.2 based on the doublet proportion distribution per cluster, thereby leading to the removal of 44 leiden clusters (resolution = 4). Additionally, single cells having greater than or equal to 2 lineage marker genes were removed to exclude potential multiplets not removed at the cluster level.

scVI Integration and cell type annotation – Batch correction and integration were performed with scVI^23^ (v0.6.8) using default parameters and normalized expression of library size 10,000. The top 8000 most variable genes were selected, based on the raw count matrices as input with sample batch and sequencing runs as covariates. The trained low-dimensional representation was used for cluster detection with the Leiden algorithm and UMAP was visualized with Scanpy^49^ v1.10.3. Clusters were manually annotated based on canonical marker genes.

Pre-processing spatial data – Space Ranger (10x Genomics, version 3.1.2) was used to demultiplex, align to mm10 reference, quantify UMIs, and generate spot-by-gene matrices and spatial coordinates for Visium libraries. Spots were filtered based on the distribution of library size, number of detected genes, and mitochondrial content to exclude low-quality spots. We normalized spot UMI counts using Scanpy v1.10.3. In order to address the variability in sequencing depth from spot-to-spot in ST assays, we normalized spots by total counts per spot using scanpy’s normalize_total function. The same Space Ranger and scanpy workflows were used for human Visium HD.

#### Differential Expression Analysis

Single-cell count matrices were pseudo-bulked at the sample level. For every sample, the raw UMI counts were summed across all cells to generate one pseudo-bulk profile for each sample. The resulting counts matrix of samples by genes was analyzed with DESeq2^1^, using a negative binomial generalized linear model, incorporating sequencing run as a batch covariate to account for technical variation. Size-factor and dispersion estimates were computed with default parameters; Wald tests for contrasts were used. Multiple testing was Benjamini–Höchberg^54^ adjusted and genes with FDR < 0.05 were considered differentially expressed.

#### GO Pathway Enrichment

Differentially expressed gene (DEG) sets were stratified by directionality, with up and down-regulated genes analyzed separately. Over-representation analysis of Gene Ontology Biological Process (GO BP)^55^ terms was performed using clusterProfiler^52^ R package (v 4.12.6).

Statistical significance of enrichment was tested with a hypergeometric test and p-values were Benjamini– Höchberg adjusted. Background (universe) gene set consisted of all genes in the dataset. Enrichment outputs were filtered at FDR < 0.05. For reducing redundancy, GO terms with similar semantics were collapsed with the simplify function in clusterProfiler (similarity cutoff = 0.7, ranking terms by adjusted p-value).

#### Cell-cell interaction using cellChat

Cell–cell interactions were inferred for every time point bin in the scRNA-seq expression dataset using CellChat^50^ (v1.6.1) in R. A cellChat object was created using the normalized gene–cell counts matrix along with cell types as input. The appropriate species-specific ligand–receptor (L-R) interaction database (CellChatDB.mouse) was used, and only L-R pairs present in the database were retained. Over-expressed genes in each cell type with default percentage of cells expressing a gene and fold-change over background thresholds were identified. Ligand–receptor interactions enriched in specific sender–receiver pairs were then detected by testing the co-expression of cognate L-R pairs.

Communication probabilities were computed based on the combined expression of ligands and receptors, with adjustment for cell population size. Statistical significance was assessed by bootstrapping with default permutations, and interactions supported by fewer than 10 cells per group were excluded. Finally, CellChat objects from each condition were merged to allow direct comparisons of the total number and overall strength of predicted interactions across time point bins, and specific L-R edges modulated across conditions.

#### Cell-type Deconvolution

We constructed a balanced single-cell reference by stratified subsampling across the early, mid and late temporal bins. We sampled without replacement within each bin to constrain the reference to ∼50,000 cells while preserving temporal composition. Feature selection retained the top 5,000 highly variable genes (Seurat v3 flavor) computed on raw counts. To deconvolute the spatial data and estimate gene expression for every cell type inside every spot DestVI^25^ v.0.1.0 was used together with the single-cell reference. Cell proportions were calculated for each spot using manual thresholds that were selected for every individual cell type based on their distribution across spots.

#### Spatial Differential Expression Analysis

For every biweekly interval, differential expression analysis was performed within every cell type against matched controls using the scRNAseq data. For each up-regulated gene, local hotspot intensity at spot i was calculated using the Getis–Ord statistic:

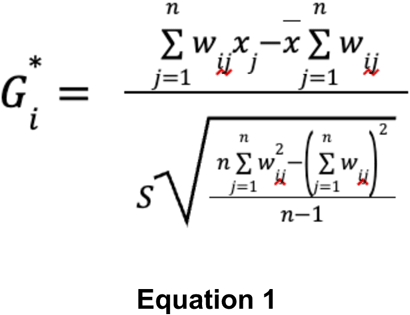

where x_j_ is the log-normalized expression of the gene at spot j, and S are the global mean and standard deviation of x across all n spots, and w_ij_ are row-standardized spatial weights. Spot centroids represented by their spatial coordinates were used to construct weights with a distance-band graph of radius 550 pixels (median ∼24 neighbors per spot), and then row-standardized. Statistical significance was assessed with default 999 random permutations, producing permutation z-scores and p-values. Multiple testing across spots was corrected with Benjamini–Höchberg and hotspots were designated at z>0 and FDR<0.05.

Hotspot calls were assembled into a binary spot-by-gene matrix H (rows: spots; columns: genes), where H_i,g_=1 if spot i is a hotspot for gene g or 0 otherwise. Pairwise spatial concordance among genes was quantified using the Matthews correlation coefficient (Φ) computed on hotspot binaries:

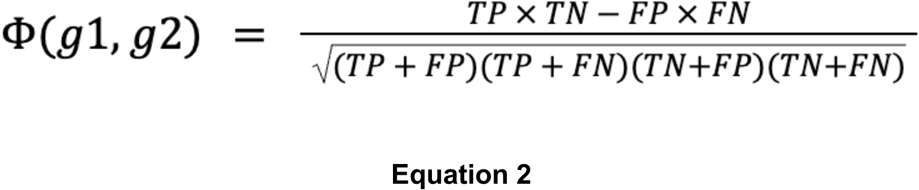

where the matrix is formed by comparing hotspot presence/absence of the two genes across spots. This yields a symmetric gene-by-gene similarity matrix with values ranging from −1 to 1. In order to sparsify the weak associations, all coefficient values below 0.5 were set to zero. This was followed by the construction of a k-nearest-neighbor graph (k = 10) in this feature space, and community detection via Louvain clustering was performed using Scanpy. The output gene clusters were taken as differentially co-localized hotspot modules.

For each Louvain cluster, over-representation analysis of Gene Ontology Biological Process terms was performed against the background of all spatially detected genes using clusterprofiler R package. Enrichment outputs were filtered at FDR < 0.05. For reducing redundancy, GO terms with similar semantics were collapsed with the simplify function in clusterProfiler package (similarity cutoff = 0.7).

#### cellSpat

Cell–cell interactions were first inferred from the single-cell RNA sequencing dataset binned using the different time points using cellchat (v1.6.1). The resulting ligand–receptor (L–R) interaction predictions were filtered to retain significant interactions based on CellChat’s communication probability score and p value < 0.05.

For each candidate interaction, the corresponding spatial transcriptomic profile of the ligand and receptor was assayed in the corresponding tissue (for example, cellchat analysis on the late bin was used for a high/invasive carcinoma tissue). We employed a non-zero median threshold to ensure sufficient signal and used it to mark “expressed” spots for every gene. In order for two genes to be considered co-localized, the ligand and receptor both needed to be above the expression threshold in the same spatial spot. Every such co-localized interaction was counted across every interaction pair.

To evaluate whether the observed co-localized interactions occurred significantly higher than expected if ligand and receptor positive spots were distributed randomly, we used a hypergeometric test. For every interaction the probability of observing at least k co-localized spots was calculated with X denoting the overlap while sampling K ligand-positive spots and M receptor-positive spots without replacement from a total of N spots as described by the survival function of the hypergeometric distribution:

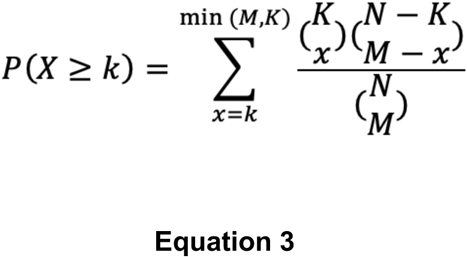

Benjamini–Höchberg false discovery rate (FDR) was used to correct for multiple testing, and the most significant interactions were ranked based on their FDR.

Co-expression for every L-R pair was calculated as the product of log-normalized expression values of the ligand and receptor across all spots. For the highest-ranked interactions, spatial “hotspots” of co-expression were identified using the Getis–Ord Gi* statistic as described in equation 1 where x_j_ is the co-expression value of the L-R pair at spot j, and S are the global mean and standard deviation of x across all n spots, and w_ij_ are row-standardized spatial weights with all parameters maintained the same as in equation 1 followed by Benjamini–Höchberg correction. Hotspots were defined by z>0 and FDR<0.05.

## REFERENCES

1. Bhat, A.A., Yousuf, P., Wani, N.A., Rizwan, A., Chauhan, S.S., Siddiqi, M.A., Bedognetti, D., El-Rifai, W., Frenneaux, M.P., Batra, S.K., et al. (2021). Tumor microenvironment: an evil nexus promoting aggressive head and neck squamous cell carcinoma and avenue for targeted therapy. Sig Transduct Target Ther 6, 12. 10.1038/s41392-020-00419-w.

2. Song, Q., Yang, Y., Jiang, D., Qin, Z., Xu, C., Wang, H., Huang, J., Chen, L., Luo, R., Zhang, X., et al. (2022). Proteomic analysis reveals key differences between squamous cell carcinomas and adenocarcinomas across multiple tissues. Nat Commun 13, 4167. 10.1038/s41467-022-31719-0.

3. Chen, J.W., and Dhahbi, J. (2021). Lung adenocarcinoma and lung squamous cell carcinoma cancer classification, biomarker identification, and gene expression analysis using overlapping feature selection methods. Sci Rep 11, 13323. 10.1038/s41598-021-92725-8.

4. Wang, C., Yu, Q., Song, T., Wang, Z., Song, L., Yang, Y., Shao, J., Li, J., Ni, Y., Chao, N., et al. (2022). The heterogeneous immune landscape between lung adenocarcinoma and squamous carcinoma revealed by single-cell RNA sequencing. Sig Transduct Target Ther 7, 289. 10.1038/s41392-022-01130-8.

5. Siegel, R.L., Miller, K.D., Fuchs, H.E., and Jemal, A. (2022). Cancer statistics, 2022. CA A Cancer J Clinicians 72, 7–33. 10.3322/caac.21708.

6. Yilmaz, E., Ismaila, N., Bauman, J.E., Dabney, R., Gan, G., Jordan, R., Kaufman, M., Kirtane, K., McBride, S.M., Old, M.O., et al. (2023). Immunotherapy and Biomarker Testing in Recurrent and Metastatic Head and Neck Cancers: ASCO Guideline. JCO 41, 1132–1146. 10.1200/JCO.22.02328.

7. Nenclares, P., Rullan, A., Tam, K., Dunn, L.A., St. John, M., and Harrington, K.J. (2022). Introducing Checkpoint Inhibitors Into the Curative Setting of Head and Neck Cancers: Lessons Learned, Future Considerations. American Society of Clinical Oncology Educational Book, 511–526. 10.1200/EDBK_351336.

8. Basil, M.C., Katzen, J., Engler, A.E., Guo, M., Herriges, M.J., Kathiriya, J.J., Windmueller, R., Ysasi, A.B., Zacharias, W.J., Chapman, H.A., et al. (2020). The Cellular and Physiological Basis for Lung Repair and Regeneration: Past, Present, and Future. Cell stem cell 26, 482–502. 10.1016/j.stem.2020.03.009.

9. Fuchs, E. (2018). Skin Stem Cells in Silence, Action, and Cancer. Stem cell reports 10, 1432–1438. 10.1016/j.stemcr.2018.04.008.

10. Jones, K.B., Furukawa, S., Marangoni, P., Ma, H., Pinkard, H., D’Urso, R., Zilionis, R., Klein, A.M., and Klein, O.D. (2019). Quantitative Clonal Analysis and Single-Cell Transcriptomics Reveal Division Kinetics, Hierarchy, and Fate of Oral Epithelial Progenitor Cells. Cell stem cell 24, 183–192 e188. 10.1016/j.stem.2018.10.015.

11. Morrisey, E.E., and Rustgi, A.K. (2018). The Lung and Esophagus: Developmental and Regenerative Overlap. Trends Cell Biol 28, 738–748. 10.1016/j.tcb.2018.04.007.

12. Feller, L., and Lemmer, J. (2012). Oral Squamous Cell Carcinoma: Epidemiology, Clinical Presentation and Treatment. JCT 03, 263–268. 10.4236/jct.2012.34037.

13. Abdalla, Z., Walsh, T., Thakker, N., and Ward, C.M. (2017). Loss of epithelial markers is an early event in oral dysplasia and is observed within the safety margin of dysplastic and T1 OSCC biopsies. PLoS ONE 12, e0187449. 10.1371/journal.pone.0187449.

14. Nokovitch, L., Maquet, C., Crampon, F., Taihi, I., Roussel, L.-M., Obongo, R., Virard, F., Fervers, B., and Deneuve, S. (2023). Oral Cavity Squamous Cell Carcinoma Risk Factors: State of the Art. JCM 12, 3264. 10.3390/jcm12093264.

15. McDowell, J.D. (2006). An Overview of Epidemiology and Common Risk Factors for Oral Squamous Cell Carcinoma. Otolaryngologic Clinics of North America 39, 277–294. 10.1016/j.otc.2005.11.012.

16. Feller, L.L., Khammissa, R.R., Kramer, B.B., and Lemmer, J.J. (2013). Oral squamous cell carcinoma in relation to field precancerisation: pathobiology. Cancer Cell International 13, 31. 10.1186/1475-2867-13-31.

17. Davis, R.J., Van Waes, C., and Allen, C.T. (2016). Overcoming barriers to effective immunotherapy: MDSCs, TAMs, and Tregs as mediators of the immunosuppressive microenvironment in head and neck cancer. Oral Oncology 58, 59–70. 10.1016/j.oraloncology.2016.05.002.

18. Lodi, G., Franchini, R., Warnakulasuriya, S., Varoni, E.M., Sardella, A., Kerr, A.R., Carrassi, A., MacDonald, L.C., and Worthington, H.V. (2016). Interventions for treating oral leukoplakia to prevent oral cancer. Cochrane Database Syst Rev 2016, CD001829. 10.1002/14651858.CD001829.pub4.

19. Mohamad, I., Glaun, M.D.E., Prabhash, K., Busheri, A., Lai, S.Y., Noronha, V., and Hosni, A. (2023). Current Treatment Strategies and Risk Stratification for Oral Carcinoma. American Society of Clinical Oncology Educational Book, e389810. 10.1200/EDBK_389810.

20. Perioperative Pembrolizumab in Head and Neck Cancer. (2025). N Engl J Med 393, 1138–1139. 10.1056/NEJMc2510882.

21. Uppaluri, R., Haddad, R.I., Tao, Y., Le Tourneau, C., Lee, N.Y., Westra, W., Chernock, R., Tahara, M., Harrington, K.J., Klochikhin, A.L., et al. (2025). Neoadjuvant and Adjuvant Pembrolizumab in Locally Advanced Head and Neck Cancer. N Engl J Med 393, 37–50. 10.1056/NEJMoa2415434.

22. Sequeira, I., Rashid, M., Tomás, I.M., Williams, M.J., Graham, T.A., Adams, D.J., Vigilante, A., and Watt, F.M. (2020). Genomic landscape and clonal architecture of mouse oral squamous cell carcinomas dictate tumour ecology. Nat Commun 11, 5671. 10.1038/s41467-020-19401-9.

23. Gayoso, A., Lopez, R., Xing, G., Boyeau, P., Valiollah Pour Amiri, V., Hong, J., Wu, K., Jayasuriya, M., Mehlman, E., Langevin, M., et al. (2022). A Python library for probabilistic analysis of single-cell omics data. Nat Biotechnol 40, 163–166. 10.1038/s41587-021-01206-w.

24. Love, M.I., Huber, W., and Anders, S. (2014). Moderated estimation of fold change and dispersion for RNA-seq data with DESeq2. Genome Biology 15, 550. 10.1186/s13059-014-0550-8.

25. Lopez, R., Li, B., Keren-Shaul, H., Boyeau, P., Kedmi, M., Pilzer, D., Jelinski, A., Yofe, I., David, E., Wagner, A., et al. (2022). DestVI identifies continuums of cell types in spatial transcriptomics data. Nat Biotechnol 40, 1360–1369. 10.1038/s41587-022-01272-8.

26. Getis, A., and Ord, J.K. (1992). The Analysis of Spatial Association by Use of Distance Statistics. Geographical Analysis 24, 189–206. 10.1111/j.1538-4632.1992.tb00261.x.

27. Matthews, B.W. (1975). Comparison of the predicted and observed secondary structure of T4 phage lysozyme. Biochimica et Biophysica Acta (BBA) - Protein Structure 405, 442–451. 10.1016/0005-2795(75)90109-9.

28. Kalluri, R. (2016). The biology and function of fibroblasts in cancer. Nat Rev Cancer 16, 582–598. 10.1038/nrc.2016.73.

29. Kim, K.-H., Won, J.H., Cheng, N., and Lau, L.F. (2018). The matricellular protein CCN1 in tissue injury repair. J. Cell Commun. Signal. 12, 273–279. 10.1007/s12079-018-0450-x.

30. Farr, L., Ghosh, S., Jiang, N., Watanabe, K., Parlak, M., Bucala, R., and Moonah, S. (2020). CD74 Signaling Links Inflammation to Intestinal Epithelial Cell Regeneration and Promotes Mucosal Healing. Cellular and Molecular Gastroenterology and Hepatology 10, 101–112. 10.1016/j.jcmgh.2020.01.009.

31. Thavayogarajah, T., Sinitski, D., El Bounkari, O., Torres-Garcia, L., Lewinsky, H., Harjung, A., Chen, H.-R., Panse, J., Vankann, L., Shachar, I., et al. (2022). CXCR4 and CD74 together enhance cell survival in response to macrophage migration-inhibitory factor in chronic lymphocytic leukemia. Experimental Hematology 115, 30–43. 10.1016/j.exphem.2022.08.005.

32. Fukuda, Y., Bustos, M.A., Cho, S.-N., Roszik, J., Ryu, S., Lopez, V.M., Burks, J.K., Lee, J.E., Grimm, E.A., Hoon, D.S.B., and Ekmekcioglu, S. (2022). Interplay between soluble CD74 and macrophage-migration inhibitory factor drives tumor growth and influences patient survival in melanoma. Cell Death Dis 13, 117. 10.1038/s41419-022-04552-y.

33. Li, R.Q., Yan, L., Zhang, L., Zhao, Y., and Lian, J. (2024). CD74 as a prognostic and M1 macrophage infiltration marker in a comprehensive pan-cancer analysis. Sci Rep 14, 8125. 10.1038/s41598-024-58899-7.

34. Jin, S., Plikus, M.V., and Nie, Q. (2024). CellChat for systematic analysis of cell–cell communication from single-cell transcriptomics. Nat Protoc, 1–40. 10.1038/s41596-024-01045-4.

35. Pellegrino, B., David, K., Rabani, S., Lampert, B., Tran, T., Doherty, E., Piecychna, M., Meza-Romero, R., Leng, L., Hershkovitz, D., et al. (2024). CD74 promotes the formation of an immunosuppressive tumor microenvironment in triple-negative breast cancer in mice by inducing the expansion of tolerogenic dendritic cells and regulatory B cells. PLoS Biol 22, e3002905. 10.1371/journal.pbio.3002905.

36. Lo, M.-C., Yip, T.-C., Ngan, K.-C., Cheng, W.-W., Law, C.-K., Chan, P.-S., Chan, K.-C., Wong, C.K.-C., Wong, R.N.-S., Lo, K.-W., et al. (2013). Role of MIF/CXCL8/CXCR2 signaling in the growth of nasopharyngeal carcinoma tumor spheres. Cancer Letters 335, 81–92. 10.1016/j.canlet.2013.01.052.

37. Lechien, J.R., Nassri, A., Kindt, N., Brown, D.N., Journe, F., and Saussez, S. (2017). Role of macrophage migration inhibitory factor in head and neck cancer and novel therapeutic targets: A systematic review. Head & Neck 39, 2573–2584. 10.1002/hed.24939.

38. Kang, Y., Zhang, Y., and Sun, Y. (2018). Macrophage migration inhibitory factor is a novel prognostic marker for human oral squamous cell carcinoma. Pathology - Research and Practice 214, 1192–1198. 10.1016/j.prp.2018.06.020.

39. Koh, H.M., and Kim, D.C. (2020). Prognostic significance of macrophage migration inhibitory factor expression in cancer patients: A systematic review and meta-analysis. Medicine 99, e21575. 10.1097/MD.0000000000021575.

40. Kindt, N., Lechien, J.R., Nonclercq, D., Laurent, G., and Saussez, S. (2014). Involvement of CD74 in head and neck squamous cell carcinomas. J Cancer Res Clin Oncol 140, 937–947. 10.1007/s00432-014-1648-9.

41. Zepeda-Nuño, J.S., Gutiérrez-Cortés, E., Hernández-Bello, J., Ángeles-Sánchez, J., De La Cruz-Mosso, U., Cruz, Á., and Muñoz-Valle, J.F. (2021). Macrophage migration inhibitory factor: A promising oncogenic serological biomarker for oral squamous cell carcinoma. Int J Immunopathol Pharmacol 35, 20587384211038417. 10.1177/20587384211038417.

42. Ma, R.-Y., Black, A., and Qian, B.-Z. (2022). Macrophage diversity in cancer revisited in the era of single-cell omics. Trends in Immunology 43, 546–563. 10.1016/j.it.2022.04.008.

43. Katkar, G., and Ghosh, P. (2023). Macrophage states: there’s a method in the madness. Trends in Immunology 44, 954–964. 10.1016/j.it.2023.10.006.

44. Zheng, G.X.Y., Terry, J.M., Belgrader, P., Ryvkin, P., Bent, Z.W., Wilson, R., Ziraldo, S.B., Wheeler, T.D., McDermott, G.P., Zhu, J., et al. (2017). Massively parallel digital transcriptional profiling of single cells. Nat Commun 8, 14049. 10.1038/ncomms14049.

45. Space Ranger | Official 10x Genomics Support 10x Genomics. https://www.10xgenomics.com/support/software/space-ranger/latest.

46. Fleming, S.J., Chaffin, M.D., Arduini, A., Akkad, A.-D., Banks, E., Marioni, J.C., Philippakis, A.A., Ellinor, P.T., and Babadi, M. (2023). Unsupervised removal of systematic background noise from droplet-based single-cell experiments using CellBender. Nat Methods 20, 1323–1335. 10.1038/s41592-023-01943-7.

47. Wolock, S.L., Lopez, R., and Klein, A.M. (2019). Scrublet: Computational Identification of Cell Doublets in Single-Cell Transcriptomic Data. Cell Syst 8, 281–291 e289. 10.1016/j.cels.2018.11.005.

48. Gayoso, A., Shor, J., Carr, A.J., Sharma, R., and Pe’er, D. (2019). GitHub: DoubletDetection (Zenodo).

49. Wolf, F.A., Angerer, P., and Theis, F.J. (2018). SCANPY: large-scale single-cell gene expression data analysis. Genome Biology 19, 15. 10.1186/s13059-017-1382-0.

50. Jin, S., Guerrero-Juarez, C.F., Zhang, L., Chang, I., Ramos, R., Kuan, C.-H., Myung, P., Plikus, M.V., and Nie, Q. (2021). Inference and analysis of cell-cell communication using CellChat. Nat Commun 12, 1088. 10.1038/s41467-021-21246-9.

51. Palla, G., Spitzer, H., Klein, M., Fischer, D., Schaar, A.C., Kuemmerle, L.B., Rybakov, S., Ibarra, I.L., Holmberg, O., Virshup, I., et al. (2022). Squidpy: a scalable framework for spatial omics analysis. Nat Methods 19, 171–178. 10.1038/s41592-021-01358-2.

52. Wu, T., Hu, E., Xu, S., Chen, M., Guo, P., Dai, Z., Feng, T., Zhou, L., Tang, W., Zhan, L., et al. (2021). clusterProfiler 4.0: A universal enrichment tool for interpreting omics data. The Innovation 2, 100141. 10.1016/j.xinn.2021.100141.

53. El-Naggar A.K., C.J.K.C., Grandis J.R., Takata T., Slootweg P.J. (2013). WHO classification of tumours of soft tissue and bone, 4th Edition (International agency for research on cancer (IARC) press).

54. Benjamini, Y., and Hochberg, Y. (1995). Controlling the False Discovery Rate: A Practical and Powerful Approach to Multiple Testing. Journal of the Royal Statistical Society Series B: Statistical Methodology 57, 289–300. 10.1111/j.2517-6161.1995.tb02031.x.

55. Ashburner, M., Ball, C.A., Blake, J.A., Botstein, D., Butler, H., Cherry, J.M., Davis, A.P., Dolinski, K., Dwight, S.S., Eppig, J.T., et al. (2000). Gene ontology: tool for the unification of biology. The Gene Ontology Consortium. Nature genetics 25, 25–29. 10.1038/75556.

