## Supplementary material for "Integrative analysis of spatiotemporal transcriptomics delineates dynamic cell states in squamous tumorigenesis": supplemental_figures.pdf

### Figure S1

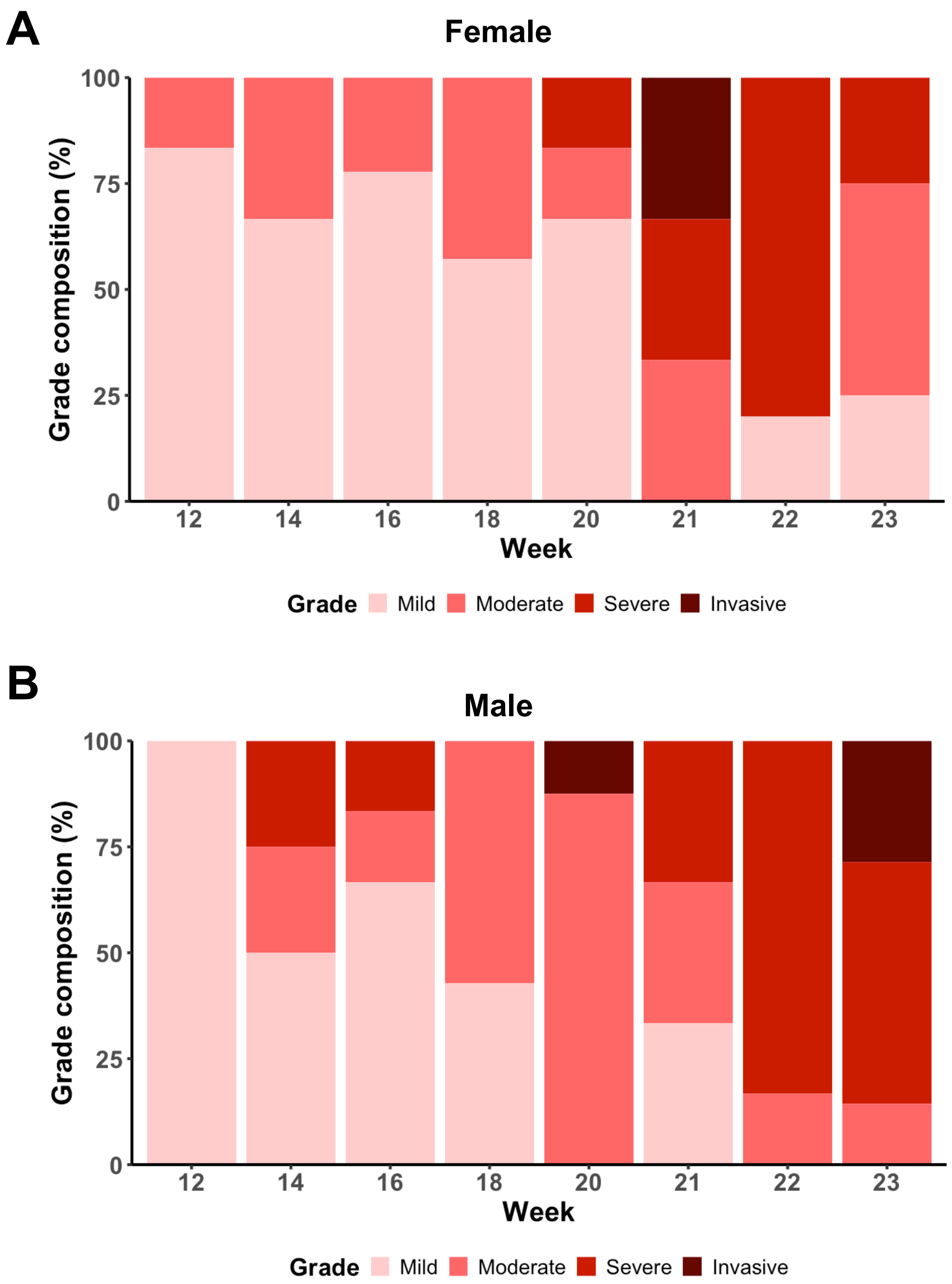


**Figure S1.** Temporal dysplasia grade distribution by sex. Bar plots show composition of dysplasia grades in **A)** female and **B)** male mice.

### Figure S2

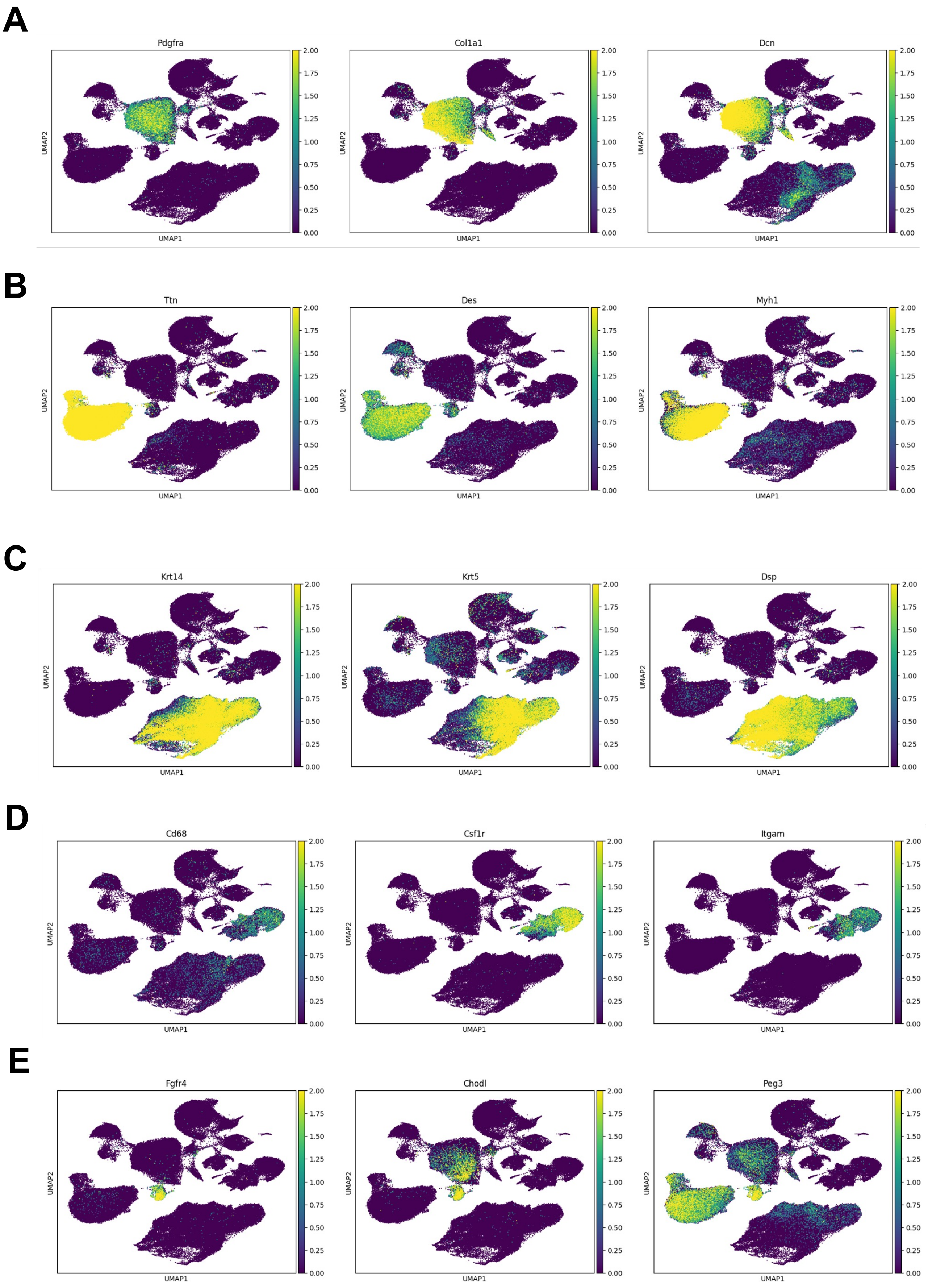


**Figure S2.** Feature plots of marker genes used for annotation cell types in scRNA-seq data: **A)** fibroblast, **B)** muscle, **C)** epithelia, **D)** macrophage, and **E)** muscle stem cells.

### Figure S3

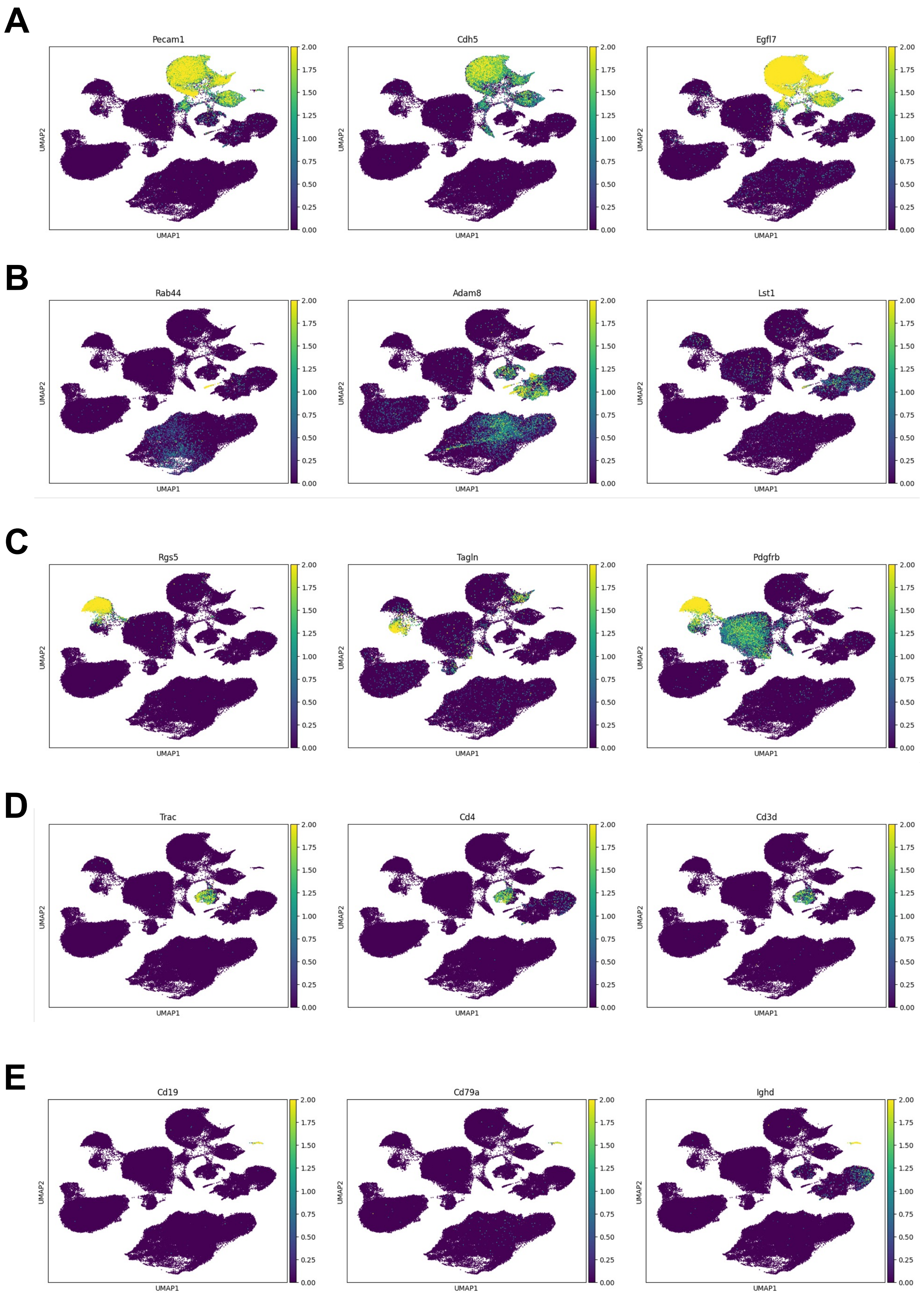


**Figure S3.** Feature plots of marker genes used for annotation cell types in scRNA-seq data: **A)** endothelial, **B)** monocytes, **C)** pericytes, **D)** T-, and **E)** B- cells.

### Figure S4

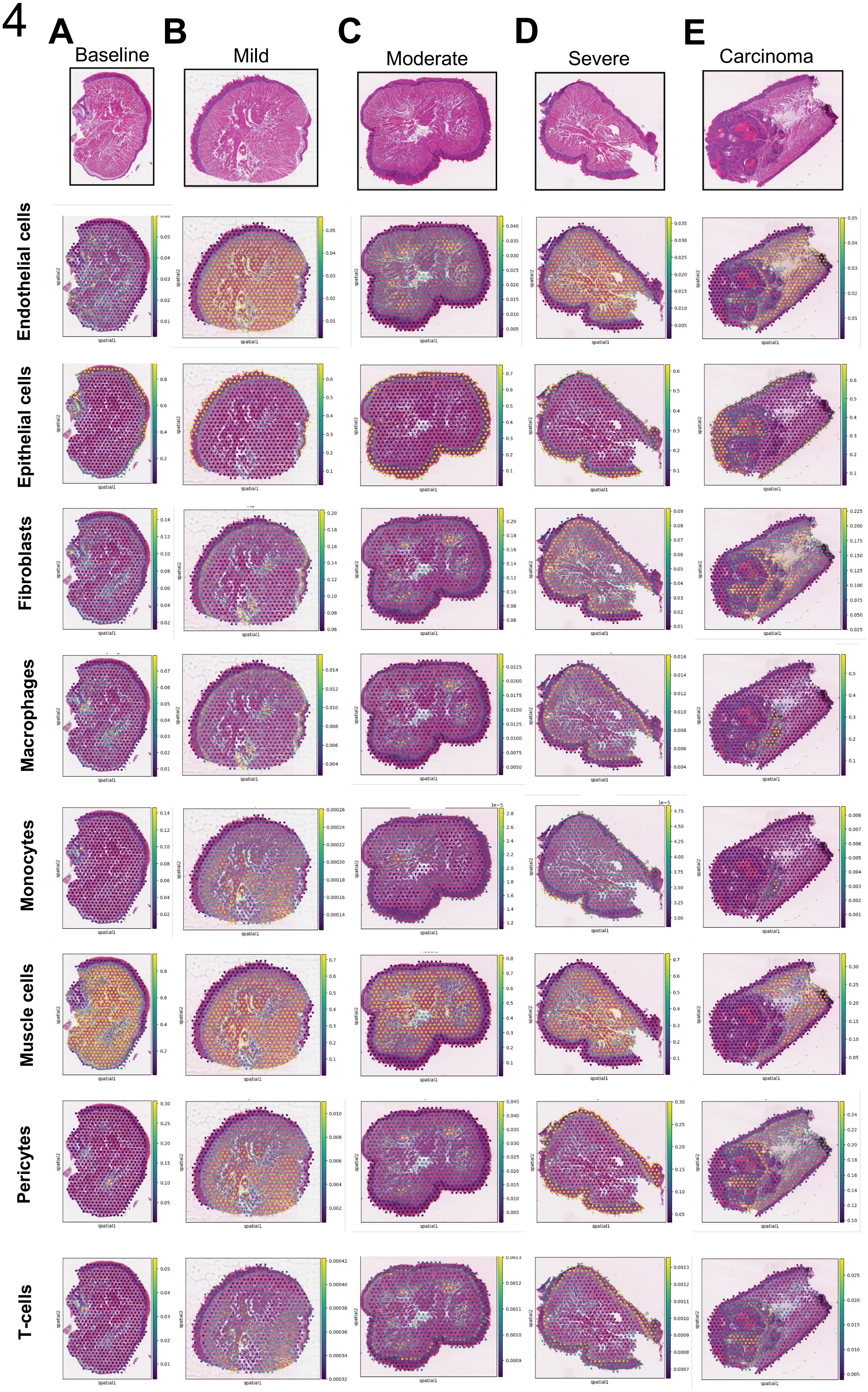


**Figure S4.** Predicted cell type proportions after deconvolution with paired scRNA-seq data in (**A**) baseline (**B**) mild (**C**) moderate and (**D**) severe dysplasia and finally (**E**) invasive carcinoma.
